# OmniCorr: An R-package for visualizing putative host-microbiota interactions using multi-omics data

**DOI:** 10.1101/2025.07.01.662509

**Authors:** Shashank Gupta, Wanxin Lai, Carl M. Kobel, Velma T. E. Aho, Arturo Vera-Ponce de León, Sabina Leanti La Rosa, Simen R. Sandve, Phillip B. Pope, Torgeir R. Hvidsten

## Abstract

Holo-omics leverages omics datasets to explore the interactions between hosts and their associated microbiomes. Although the generation of omics data from matching host and microbiome samples is steadily increasing, there remains a scarcity of computational tools capable of integrating and visualizing this data to facilitate the interpretation and prediction of host-microbiota interactions. We present **OmniCorr**, an R package designed to: (1) manage the complexity of omics data by clustering similar observations into modules, (2) visualize correlations of these modules across different omics layers, host-microbiota interfaces, and meta-data, and (3) identify statistically significant associations indicative of putative host-microbiota interactions. OmniCorr’s utility is demonstrated using datasets from two systems: (i) Atlantic salmon, integrating host transcriptomics with metagenomics and metatranscriptomics to explore dietary impacts, and (ii) cattle, combining host proteomics with metaproteomics and metabolomics to investigate methane emission variability.

**Availability and Implementation:** OmniCorr is freely available at: https://github.com/shashank-KU/OmniCorr.

## Introduction

The holobiont, comprising a host organism and its associated microbiota, represents a complex biological system with intricate interactions that significantly influence health and disease states (Robinson and Cameron, 2020; Bordenstein and Theis, 2015). These host-microbiome interactions play crucial roles in various physiological processes, including metabolism, immune function, and overall well-being (Bordenstein and Theis, 2015; Rowland *et al*., 2018). Multi-omics is increasingly being used for molecular profiling of matching samples from host tissues and gut microbiota collected, for example, during feeding trials designed to modulate the gut microbiota or across individual animals with variations in physiological traits believed to be affected by the gut microbiota (Muller *et al*., 2024; Kwoji *et al*., 2023).

The aim of such holo-omics studies is to identify putative interactions between molecular markers in the host and the microbiota that could confer an effect on the host (Nyholm *et al*., 2020). The multi-omics datasets generated from holo-omics studies present a number of data analysis challenges, one of which is the large number of variables (genes, proteins, metabolites, microbial species: ranging from thousands to millions) compared to the relatively small number of samples (ranging from tens to a few hundreds) (Zhao *et al*., 2019).

The task of discerning true host-microbiota associations from those driven by random variability or single outlier features remains challenging, although several computational methods have been proposed (Kobel *et al*., 2024).

In contrast to many existing multi-omics tools that rely on latent factor modeling and other statistical frameworks (e.g., mixOmics, MOFA) (Rohart et al., 2017; Argelaguet et al., 2020), OmniCorr is explicitly designed as an exploratory tool that aims to generate biologically interpretable hypotheses by directly correlating hub features across omics layers and across the host-microbiota boundary. By offering flexible and customizable visualizations, OmniCorr complements—but does not replace—more complex integrative frameworks. This design is particularly advantageous for studies with limited sample sizes and interest in interpretable host-microbiota interactions.

Here we describe an easy-to-use R package, OmniCorr, designed to seamlessly integrate multiple omics datasets from a host and its microbiota. OmniCorr simplifies data visualization, enabling intuitive exploration of complex datasets. Compared to existing methods (Munk *et al*., 2024), it is more flexible, user-friendly, and efficient, offering a streamlined workflow that reduces the learning curve for non-experts.

### The OmniCorr package

The OmniCorr method approaches the challenge of predicting host-microbiota interactions from omics data by first reducing the dimensionality of the omics data and then by visualizing correlations across omics-layers, host-microbiota boundaries and between omics data and meta-data.

The package can be installed with the command:

~~~
devtools::install_github(“shashank-KU/OmniCorr”
~~~

and consists of the following steps:

**Step 0: Reduce dimensionality of omics data**: This step allows the user to choose their preferred method for clustering or projection. We recommend the robust inference of network modules as implemented in the R package Weighted Gene Co-expression Network Analysis (WGCNA) (Langfelder and Horvath, 2008). Here, the function `blockwiseModules` identifies modules and the function `chooseTopHubInEachModule` extracts the most connected gene, protein or metabolite (hub) from each of these modules. This hub selection is based on intramodular connectivity, as implemented in the WGCNA package, where it has been shown to produce biologically meaningful and robust module representatives (Langfelder and Horvath, 2008). Although other representations such as eigengenes or centroids may be used, the default selection of the top hub is grounded in the reproducibility and interpretability of WGCNA-defined hubs across diverse datasets. The output from step 0 is thus a table for each omics layer with samples as rows (these samples must match across all omics layers and species/kingdom) and hubs as columns. Alternatives to hubs are e.g. centroids or the first principal component (eigengene) of each cluster/module.

**Step 1: Perform hierarchical clustering of the central omics layer using WGCNA**: For the visualization, the user selects one central omics layer where the profiles of the hubs across samples will be displayed as a heatmap. The other omics layers will appear in the visualization only as hub-hub-correlations. The hubs in the chosen omics layer are ordered using hierarchical clustering (`hclust`-function).

**Step 2: Generate a heatmap of the central omics layer with dendrogram**: Given the hierarchical clustering from Step 1 (i.e. the dendrogram), a heatmap is generated for the central omics layer using the `pheatmap`-function.

**Step 3: Calculate correlations between omics data**: The OmniCorr function `calculate_correlations` computes correlations between the modules (represented by one hub per module) in the central omics layer and the modules of other omics layers. If the centrepiece of the visualization is from the host, the other layers are typically from the microbiota. The correlations indicate to what degree modules in the central layer co-vary across the samples with the modules of each of the other layers. Significance for the correlations can also be computed and added to downstream correlation heatmaps. The output is a module-module correlation matrix per omics layer. To provide flexibility and transparency in statistical analysis, the `calculate_correlations()` function allows users to specify the correlation method (“pearson”, “spearman”, or “kendall”), handle missing data using various approaches (e.g., “all.obs”, “pairwise.complete.obs”), and choose the method for adjusting p-values for multiple testing (e.g., “fdr”, “bonferroni”). The function computes correlation coefficients between all variable pairs from the two input omics layers, tests for statistical significance, and returns matrices of correlation values, raw p-values, adjusted p-values, and annotated significance levels. Significance can be reported as asterisks (e.g., *, **, ***), exact p-values, or raw correlation values, depending on user preference. This enables users to tailor the analysis to the context of their specific dataset and to interpret correlations with statistical confidence.

As OmniCorr is designed to be agnostic to specific omics types, it does not impose specific preprocessing steps designed for e.g. sparse or compositional data. Instead, users are expected to preprocess these layers according to best practices, e.g., filtering low-prevalence features, applying centered log-ratio (CLR) transformation, or normalizing via cumulative sum scaling (CSS). This modularity allows users to tailor preprocessing to the statistical properties of their data before applying OmniCorr for integrative correlation analysis.

**Step 4: Generate a heatmap of the correlations between the central layer and other layers:** Heatmaps for correlations from Step 3 are generated using the `pheatmap`-function.

**Step 5: Combine the heatmap of transcriptomics data and the heatmap of correlations**: Heatmaps from Step 2 and Step 4 are combined using the `plot_grid`-function.

**Step 6: Integrate the meta-data heatmap:** Finally, the OmniCorr function `calculate_correlations` is used to compute correlations between the modules in the central omics layer and meta-data. The correlations are represented as a heatmap, annotated with significant associations, and the meta-data correlation heatmap is combined with the heatmaps from Step 5 to display an integrated overview of correlations across omics layers and host-microbiota boundaries.

## Additional analysis

Other tools such as the R package `clusterProfiler` (Yu *et al*., 2012) can be used to perform Gene Ontology (GO) and pathway enrichment analysis of the modules. Enriched functions, pathways or bacterial phyla can be used to name modules in the visualization in order to enhance interpretation. Enrichment analysis methods can vary depending on the omics layer (e.g., microbial species, genes, proteins), OmniCorr outputs are structured for compatibility with standard enrichment packages.

### Case study 1: Feed-host-microbiome interaction in Atlantic salmon

Microbiome-directed dietary interventions, such as microbiota-directed fibres (MDFs), have been effective in promoting beneficial gut microbes in several animal production systems. However, there are knowledge gaps related to the specific metabolic influence that such feed supplements exert in their host and/or their microbiota. To investigate the effects of dietary supplementation on Atlantic salmon (*Salmo salar*), Gupta et al. (Gupta *et al*., 2024) conducted a feeding trial designed to assess individualized responses to an industry-standard 0.2% inclusion of one acetylated β-galactoglucomannan (MN3) and two different types of α-mannans (MC1 and MC2), in comparison to a standard diet (CTR). This trial lasted for 110 days, transitioning from freshwater to seawater, with samples collected at four distinct time points (T0, T1, T2, and T3). The omics analysis included host transcriptomics from the hindgut (69,389 genes), microbial community characterization through 16S rRNA gene amplicon sequencing (6,468 Amplicon Sequencing Variants - ASVs -identified), and metatranscriptomics (117,261 genes). Additionally, the collected meta-data comprised measurements of fish weight, length, gutted weight, condition factor, hepatosomatic index, cardio somatic index, welfare indicators and organ integrity (Gupta *et al*., 2024). In the present study, we applied OmniCorr to this dataset, while excluding the samples from the T0 sampling time point (as no experimental diet was provided during that period). We reduced the dimensionality of omics data and identified 24 host modules, 6 microbial modules, and 14 metatranscriptome modules. Several host and microbiota modules were correlated with sampling time points, whereas no effect was detected for the feed additive mannan (**Figure 1A**), in agreement with our previous results (Gupta *et al*., 2024).

**Figure 1.**
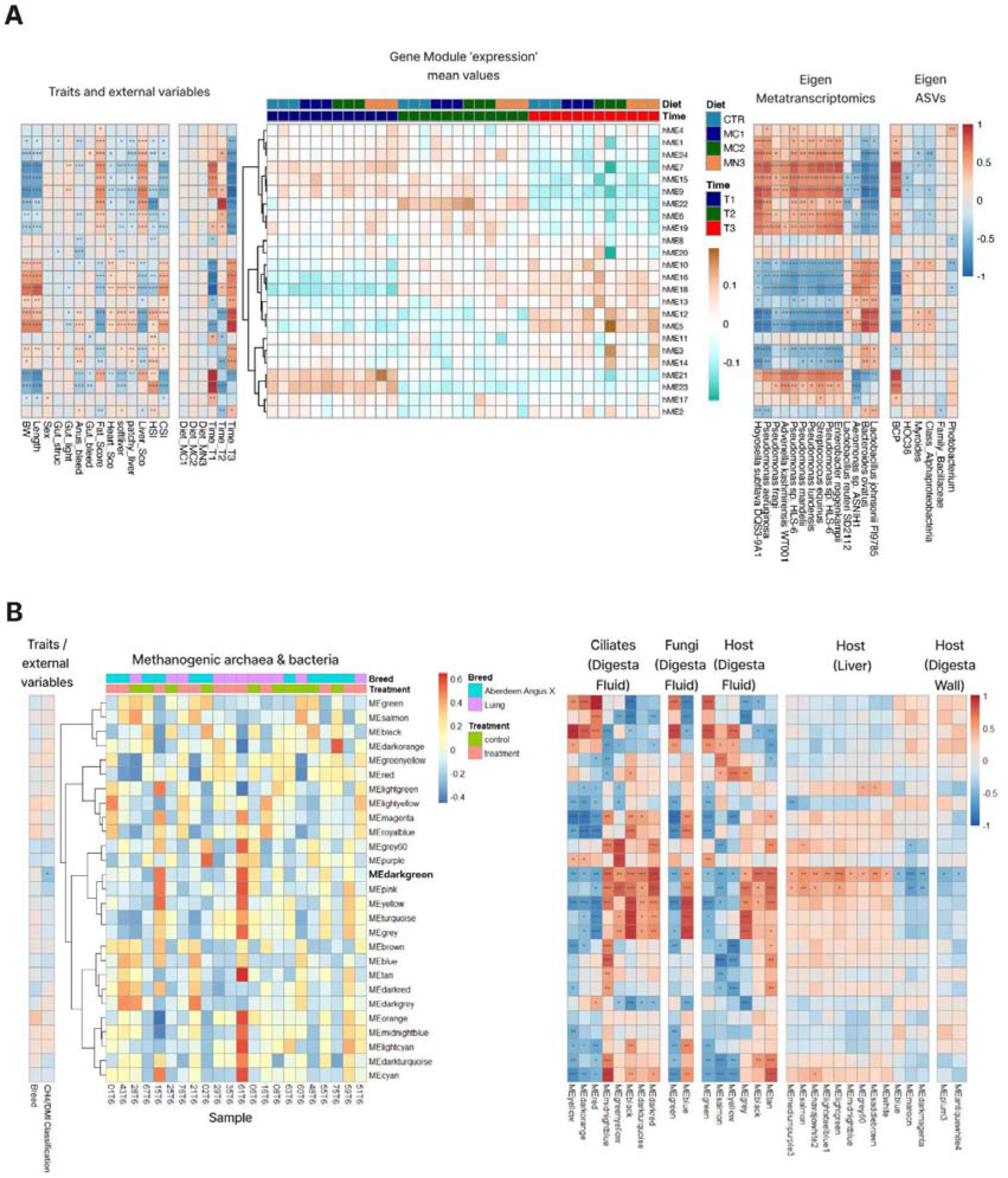
Two case studies applying OmniCorr to multi-omics data from (A) Atlantic salmon and (B) cattle. The central heatmap visualizes the central omics layer with columns corresponding to samples and rows corresponding to modules (represented by the eigengene). Correlations between the central omics layer and metadata are visualized to the left, with stars indicating significant correlations to modules in various omics layers are visualized to the right. Binary classification of methane emission efficiency (CH4 per dry matter intake, CH^_^/DMI) into ‘low’ and ‘high’ categories was the only metadata variable significantly correlated with the module eigengene (MEdarkgreen).

Host modules hME9 and hME5 demonstrated strong correlations with the water environment (fresh *vs* salt). Correlations between the host omics data and water environment changes is expected in this dataset as Atlantic salmon undergoes complex preparatory physiological changes to an adult life in seawater and experience additional acute physiological responses when salinity levels increase (Harvey *et al*., 2024). Furthermore, the host module (hME9) was strongly and significantly positively correlated with ASVs from the *Burkholderia-Caballeronia-Paraburkholderia* (BCP) group, suggesting a close relationship between the host’s molecular responses and specific microbial communities. This correlation highlights potential host-microbe interactions, where the identified microbial taxa may play supportive roles in facilitating host adaptation during the transition to seawater. In the BCP group, we identified genera such as *Pseudomonas*, which were also detected in the metatranscriptomics data and exhibited strong positive correlations with host module (hME9) (**Figure 1A**).

Moreover, manual inspection of gene expression by BCP taxa in Gupta et al. (Gupta *et al*., 2024) highlighted the metabolic influence this group can exert towards digestion and generation of energy-yielding nutrients.

### Case study 2: Methane emission in different cattle breeds

Livestock significantly contribute to global greenhouse gas emissions, particularly methane (Blaustein-Rejto *et al*., 2023), which is produced during digestion in the rumen where complex microbial communities reside (Grossi *et al*., 2019). Methane is a by-product of methanogenesis, a process driven by methanogenic archaea. While bacteria do not directly participate in methanogenesis, they break down complex carbohydrates, producing by-products such as hydrogen, which fuel other microbes and support methanogens. Understanding the interactions between the host and its microbiome can therefore help reduce emissions by pinpointing microbial activities that can be targeted without affecting livestock productivity. In this study, we integrated (meta)proteomics data from multiple layers of the rumen ecosystem: methanogenic archaea, bacteria, ciliate protozoa, and fungi (all from digesta fluid samples), as well as host tissues (digesta fluid, liver, and rumen wall). Twenty-two samples were collected from high- and low-methane emitting cattle (Luing and Aberdeen Angus X), which were managed under identical conditions but defined by emissions above or below 24 CH□ [g/kg DMI] (measured in respiration chambers). By focusing on the methanogenic Archaea and Bacteria (ArcBac) as the central omic layer, we aimed to explore interspecies interactions that influence methane emissions, complementing proteomics insights from host tissues (Kobel *et al*., 2024).

We applied OmniCorr to this dataset and identified 27 modules from ArcBac, 31 from ciliates (6,473 proteins), and 7 from fungi (82 proteins), respectively. In the host, we found 15 modules (222 proteins) from digesta fluid, 68 modules (2,389 proteins) from liver, and 92 modules (2,619 proteins) from the digesta wall (**Figure 1B**). Notably, the darkgreen module from ArcBac showed a significant correlation with methane emission, while 8, 2, 6, 12, and 2 modules from ciliates, fungi, host digesta fluid, liver, and rumen wall, respectively, were also correlated with this module. The darkgreen module, enriched for enzymes involved in streptomycin biosynthesis, showed 22 proteins with lower abundance in high methane emitters compared to low emitters (**Supplementary Fig. 1 and 2**). How elevated streptomycin production could impact the ruminal metabolic networks that give rise to methanogenesis is a tantalizing question but is supported by evidence that antibiotic use (e.g. monensin) has been used to effectively reduce methane levels in cattle (Appuhamy *et al*., 2013). Moreover, strong correlations between this module and ciliates in terms of organismal systems and metabolism (KEGG pathways, **Supplementary Fig. 1**) suggests that ciliates modulate their archaeal and bacterial partners, coordinating complex symbiotic interactions within metabolic networks (Kobel *et al*., 2024; Rossi *et al*., 2019). Cross-domain interaction between the ArcBac and fungi was observed for the blue and yellow modules, which were enriched in nitrogen metabolism. This finding suggests the influence of fungi on shaping the microbial community by altering the availability of ammonia or nitrates and hydrogen, which has an indirect effect on methanogens (Yang *et al*., 2016). Importantly, some host modules also demonstrated significant correlations with both methane emission levels and the darkgreen module from ArcBac. The black module, enriched for G protein-coupled receptors (GPCRs), opens future hypotheses to test if these proteins in the cattle gut could contribute to physiological functions that impact digestion and metabolic efficiency. Additionally, the enrichment of pathways (black module) related to serotonergic synapse, neuroactive ligand-receptor interactions, and axon regeneration are in line with previous findings reporting that neurotransmitters in the enteric nervous system modulate gut mobility, secretion, and the microbial community in response to stress signaling (Layunta *et al*., 2021; Stepniewski *et al*., 2021). Finally, the midnightblue module from the cattle liver was enriched for pathways highly relevant to immune and inflammatory responses: phagosome, peroxisome, endocytosis, and neutrophil extracellular trap formation. These modules contained proteins with higher abundance in low emission cattle compared to high emission cattle (Li *et al*., 2023; Ma *et al*., 2024).

## Discussion

We showcase two examples of vastly different animal host-gut microbiome systems, where OmniCorr can unveil possible host-microbiota interactions that could be tested in future analyses. Although the Atlantic salmon feeding trial revealed no effect of adding 0.2% mannan to the diet (Gupta *et al*., 2024), the analysis highlighted several other potentially interesting correlations across the host-microbiota axis.

The cattle-emission analysis based on the data from Kobel et al. (Kobel *et al*., 2024) revealed a network of 22 key modules across microbial- and host-derived proteins linked to methane emission. Our results are in line with previous findings that have shown that “high methane emission” rumina prioritize energy harvesting and are dominated by methanogens and their associated fermentative bacteria (Khairunisa *et al*., 2023; Pereira *et al*., 2022; Shabat *et al*., 2016; Wang *et al*., 2017). Meanwhile, the enriched pathways identified from host-associated modules were related to neurotransmission, inflammatory responses, and cellular signalling, proposing a role of the host in coordinating microbial activity and digestion, which contributes to altered metabolic efficiency. Understanding these interactions may offer insights into how systemic physiological responses, influenced by both host and microbial factors, contribute to methane emission.

To address the robustness of associations identified by OmniCorr, we evaluated their biological plausibility in both case studies. For example, in the Atlantic salmon dataset, the correlation between host module hME9 and the *Burkholderia-Caballeronia-Paraburkholderia* group is consistent with known roles of these taxa in nutrient metabolism and adaptation to saltwater environments. Similarly, in cattle, the ArcBac darkgreen module correlated with methane emission and was enriched for streptomycin biosynthesis — a pathway previously implicated in methane suppression via antibiotic use (Appuhamy et al., 2013). These findings suggest that OmniCorr captures biologically meaningful signals.”

We recognize that tools such as mixOmics and MOFA have advanced the field of multi-omics integration by providing latent variable models capable of explaining shared variance across datasets. However, these methods may be less accessible to users focused on transparent interpretation and hypothesis generation from limited sample sizes. OmniCorr is designed to complement such tools by providing a user-friendly, statistically transparent workflow for visualizing significant cross-omics correlations. It serves as a platform to pinpoint interpretable associations that warrant further validation and explicitly offers solutions to analyze host-microbiota systems.

In conclusion, while omics layers can be analyzed separately with respect to e.g. dietary effects, we show that, using the OmniCorr package, new insight can be gained from an integrated analysis of the host and its associated microbiome. Our computational approach enables studying the system as holobiont, which takes into account the interplay between the host and its associated microbiome.

## Acknowledgements

This work was supported by the Research Council of Norway (project no. 300846), the Swedish Research Council Formas (grant no. 2019-02336) and the European Union’s Horizon 2020 research and innovation program under the ERA-Net Cofund project BlueBio (grant agreement no. 311913). The Orion High Performance Computing Center at the Norwegian University of Life Sciences and Sigma2 - the National Infrastructure for High Performance Computing and Data Storage in Norway are acknowledged for providing computational resources that have contributed to meta-omics analyses described in this study. The authors thank Elixir-Norway (NFR project no. 322392) for bioinformatics and data management related services.

