## Supplementary Fig. 1 and 2 for "OmniCorr: An R-package for visualizing putative host-microbiota interactions using multi-omics data"

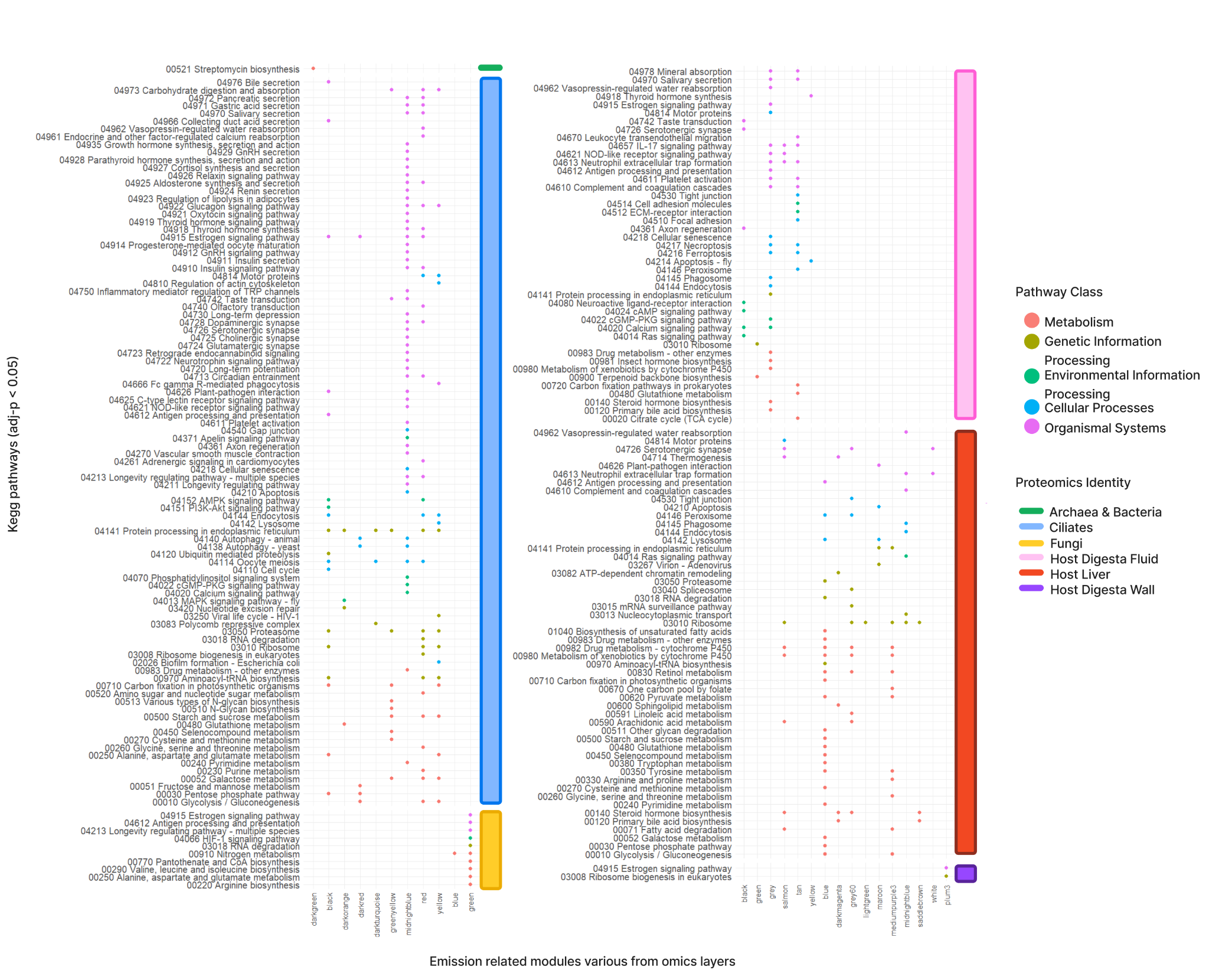

*Supplementary Figure 1: Enriched KEGG pathway modules (adj. P-value < 0.05) identified across various omics layers.*

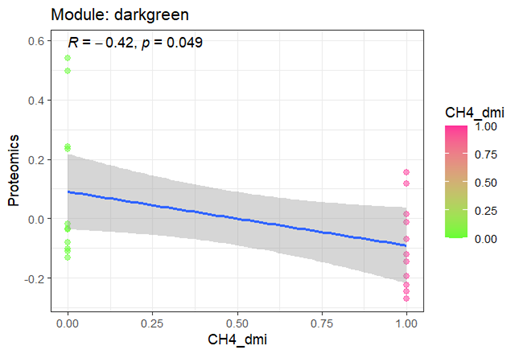

*Supplementary Figure 2: 22 proteins from the darkgreen modules with higher abundance in low methane emission cattles than high methane emission cattles.*
